# Intensity-dependent corticospinal facilitation by repetitive peripheral magnetic stimulation: Evidence for a major contribution of group I afferents

**DOI:** 10.64898/2025.12.29.696815

**Authors:** Kaito Yoshida, Mitsuhiro Nito, Dai Miyazaki, Ayu Omiya, Kanau Shitara, Tadaki Koseki, Daisuke Kudo, Nariyuki Mura, Hiromi Fujii, Tomofumi Yamaguchi

## Abstract

**Background:** Repetitive peripheral magnetic stimulation (PMS) is increasingly used in neurorehabilitation, yet the optimal stimulation intensity for inducing corticospinal facilitation and the underlying afferent mechanisms remain unclear. We investigated the intensity-dependent effects of repetitive PMS on corticospinal excitability and single motor unit responses, and tested group I afferent contribution.

**Methods:** Healthy participants received repetitive PMS (25 Hz; 2-s ON/2-s OFF) over the extensor carpi radialis (ECR) in a crossover design at 0.9× motor threshold (MT), 1.2×MT, and high intensity sufficient to induce maximal wrist dorsiflexion (mean 1.8×MT). Motor-evoked potentials (MEPs) elicited by transcranial magnetic stimulation were recorded from the ECR and flexor carpi radialis (FCR) before and during the intervention (total 15 min). The lasting effects were assessed after 9 min of high-intensity PMS for 50 min. To examine group I afferent contribution, the same high-intensity protocol was applied during upper-arm ischemia after reducing the ECR H-reflex to <10% of baseline. Sensory–motor input characteristics across stimulation intensities were compared using post-stimulus time histograms of ECR single motor unit firings during weak voluntary contraction.

**Results:** High-intensity PMS significantly increased ECR MEPs after 9 min of intervention, whereas 1.2×MT of PMS required 15 min to induce a marked effect. PMS at 0.9×MT did not induce significant MEP changes. Across all intensities, the FCR MEPs remained unaltered. ECR MEPs remained markedly elevated for up to 30 min after 9 min of high-intensity PMS. In contrast, PMS delivered during ischemia produced no MEP enhancement. The motor unit analysis revealed that suprathreshold PMS elicited an early peak in firing probability—consistent with monosynaptic Ia excitation—whose amplitude increased with stimulation intensity, whereas PMS at 0.9×MT produced no discernible peak.

**Conclusions:** Repetitive PMS above MT facilitates corticospinal excitability in an intensity-dependent manner. Facilitation was abolished during ischemia. Together with the presence of a short-latency peak in motor unit firing via a monosynaptic pathway, this finding supports a major contribution of large-diameter muscle afferents, with a substantial Ia component, to PMS-induced corticospinal facilitation.

## BACKGROUND

The corticospinal tract is the primary pathway that mediates voluntary motor control in humans (1, 2). Transcranial magnetic stimulation (TMS)–elicited motor-evoked potentials (MEPs) provide a noninvasive measure of corticospinal excitability (CSE) (3). CSE changes have been closely linked to motor impairment and recovery after central nervous system (CNS) lesions (4), and to motor performance improvement (5, 6). These associations underscore CSE’s central role in motor function and its modulation as a potential therapeutic target.

Repetitive peripheral magnetic stimulation (PMS) has emerged as a potential adjuvant therapy in physical rehabilitation and is increasingly used to promote motor recovery after CNS lesions (7–11). Accumulating evidence suggests that repetitive PMS facilitates motor recovery by enhancing cortical excitability (7, 12, 13). However, despite these promising findings, the optimal stimulation intensity for inducing neuroplastic changes remains poorly understood. Previous studies have applied various PMS intensities, including levels inducing palpable muscle contractions or small joint movements (7, 9, 10, 12–16) and levels producing maximal joint movements (17, 18). Others have prescribed PMS as a fixed percentage of the maximal stimulator output without calibrating intensity to the target muscle response (19–22). Overall, despite evidence supporting contraction-inducing PMS for enhanced motor recovery after CNS lesions, systematic characterization of its physiological, stimulation intensity–dependent effects on CSE remains unestablished.

PMS physiological effects are thought to result primarily from the activation of low-threshold afferents, inducing Ia afferents from muscle spindles, Ib afferents from Golgi tendon organs, and cutaneous afferents (23–25). Studies using peripheral electrical stimulation show that suprathreshold, contraction-inducing stimulation increases CSE, whereas subthreshold stimulation can reduce it, likely through preferential activation of inhibitory afferent pathways (26–28). Despite expectations of similar mechanisms with PMS, afferent and efferent fiber recruitment patterns differ from those observed with peripheral electrical stimulation. In general, electrical stimulation with increasing strength elicits action potentials in the largest fibers first, as they have the lowest electrical resistance, followed by progressively smaller fibers (23). In contrast, peripheral nerve magnetic stimulation requires a higher excitation threshold for sensory fibers than for α-motor fibers, and H-reflexes are not elicited at intensities below the motor threshold (MT) (29). Moreover, PMS generates eddy currents that directly stimulate deeper tissues without passing through the skin, suggesting reduced recruitment of cutaneous afferents (8). Collectively, these properties imply that subthreshold PMS may fail to activate low-threshold muscle afferents and may therefore be insufficient to enhance CSE. Clarifying subthreshold PMS effects will provide insight into the stimulation intensity–dependent mechanisms of PMS-induced neuroplasticity and help optimize stimulation parameters for diverse clinical applications. In addition, PMS-induced CSE modulation lacks direct afferent pathway characterization. In particular, the extent of group I afferent—especially muscle spindle group Ia afferent—contributions to repetitive PMS–induced CSE facilitation remains unclear.

Therefore, this study investigated the effects of different repetitive PMS intensities applied to the wrist extensor muscles on CSE. Group I afferent contributions were assessed using ischemic nerve block and motor unit (MU) firing–based afferent input estimates. We hypothesized that 1) suprathreshold PMS would trigger intensity-dependent CSE enhancement and within a shorter intervention period than subthreshold PMS, and 2) these effects would be mediated primarily by large-diameter group I afferents, with findings consistent with a major contribution from Ia afferents. Four experiments were conducted: Experiment 1 tested intensity-dependent time courses of MEPs; Experiment 2 examined persistence after a short high-intensity protocol; Experiment 3 probed the necessity of large-diameter afferent input using ischemia after marked H-reflex suppression; and Experiment 4 quantified afferent-driven motoneuron facilitation using single MU recordings.

## METHODS

### Participants

The study included 34 healthy volunteers, whose participation was distributed across four experiments (20 females, aged 18–34 years, 23 ± 4 years, mean ± standard deviation [SD]). The number of participants for Experiments 1, 2, 3, and 4 was 19 (10 females, 24.0 ± 4.2 years), 17 (9 females, 22.9 ± 4.7 years), 5 (all males, 27.4 ± 4.2 years), and 8 (3 females, 25.6 ± 4.1 years), respectively. Notably, six individuals participated in multiple experiments. Specifically, three individuals completed all four experiments. One individual completed Experiments 1, 2, and 4, while two individuals completed Experiments 2, 3, and 4. None of the participants had a history of neurological disease or received any medication affecting the CNS. All participants except two were considered right-handed based on Chapman’s dominant hand test (30). The experimental procedures were approved by the Ethics Committee of Yamagata Prefectural University of Health Science (approval number: 2308-15) and followed the Declaration of Helsinki. Before participating in this study, all participants signed written informed consent forms for the experimental procedures.

### Experimental setup

#### Electromyographic recordings

Surface electromyographic (EMG) signals were recorded with paired 1.0-cm diameter Ag/AgCl disk electrodes. The electrodes (1.5 cm interelectrode distance) were secured to the skin overlying the right extensor carpi radialis (ECR) and flexor carpi radialis (FCR) muscles. The EMG signals were amplified, bandpass filtered (15–1,000 Hz), and sampled at 2,000 Hz for offline analysis (Micro 1401 with Signal software, Cambridge Electronic Design, Cambridge, UK).

MU discharges were recorded with a pair of needle electrodes (Seirin acupuncture needle, 0.16□mm diameter, Seirin Kasei, Shizuoka, Japan) inserted into the ECR muscle belly. EMG signals were amplified and processed with a low-cut filter (50□Hz). Audiovisual EMG potential feedback was provided to help participants maintain stable single MU activity. Single MU discrimination was carefully defined using the upper and lower amplitude discrimination thresholds of the recorded potential. The EMG potential shape was displayed on an oscilloscope (TDS210, Tektronix, Tokyo, Japan) throughout the experiment. The MU discharges were converted into standard pulses and used for offline analysis (see *Experiment 4*).

#### Peripheral magnetic stimulation

PMS was applied over the right ECR muscle to stimulate the radial nerve innervating the ECR, as described previously (13). The forearm was fixed in a pronation position with the fingers free. A biphasic pulse of PMS was delivered using a figure-eight coil (Cool-B65; outer diameter 75 mm) connected to a MagPro R30 (MagVenture A/S, Denmark). The coil was placed on the skin overlaying the right ECR muscle and positioned perpendicular to the forearm. The stimulus intensity was expressed in multiples of the MT of the direct motor response (M-wave). MT was defined as the minimal stimulus intensity required to induce an M-wave ≥ 50 μV (peak-to-peak amplitude) by a single-pulse stimulus in at least three of five consecutive trials. The ECR muscle contraction was confirmed by palpation, with the stimulus intensity well above the MT.

#### Transcranial magnetic stimulation

TMS was applied over the left primary motor cortex using a figure-eight coil (loop diameter 70 mm) connected to Magstim 200 (Magstim Company, Whitland, Dyfed, United Kingdom). We determined the optimal positioning to elicit MEPs in the ECR muscle at rest (hot spot) by moving the coil with the handle pointing backward and 45° away from the midline. The hot spot was defined as the region where the largest MEP in the ECR muscle could be evoked with minimal stimulus intensity (Lotze et al., 2003). Resting MT was defined as the minimal stimulus intensity required to induce MEPs of ≥ 50 μV (peak-to-peak amplitude) in at least three of five consecutive trials in the relaxed muscle (5). The TMS intensity was set at 120% of the resting MT to measure MEPs as a CSE indicator. A total of 11 MEPs was recorded in the resting condition. Each peak-to-peak amplitude was measured and averaged, and the mean value among the participants was calculated for further analysis.

#### Electrical peripheral nerve stimulation

Rectangular electrical pulses (1.0 ms) were percutaneously delivered to the radial nerve trunk using bipolar surface electrodes (1.0 cm diameter, 1.5 cm interelectrode distance) positioned along the nerve trajectory at the arm’s lateral intermuscular septum. The stimulus electrode was connected to a constant-current stimulator (DS8R, Digitimer, Welwyn Garden City, UK). The maximum direct motor response (M-max) was measured by supramaximal electrical stimulation (at an intensity of 120% to induce M-max). The inter-stimulus interval was 4–5 s.

### Experimental procedures

Participants were comfortably seated, with the examined right arm supported on an armrest; the shoulder was slightly flexed (≈10°), followed by elbow flexion (≈90°), and forearm pronation.

#### Experiment 1: Intensity-dependent effects of repetitive PMS on MEPs, M-waves

Repetitive PMS was delivered in a 2 s ON and 2 s OFF cycle, according to a previous report (13). The stimulation frequency was set at 25 Hz for the following reasons: 1) the stimulation frequency of repetitive PMS was determined by afferent input from muscle spindles (Ia afferents) of the wrist extensor muscles during voluntary movement (31); 2) our previous study demonstrated that 25 Hz and 50 Hz repetitive PMS produced comparable increases in MEPs (13); and 3) in our preliminary experiment, application of 50 Hz repetitive PMS at higher intensities caused stimulation-coil heating, which prevented some participants from completing a 15-min intervention.

The effects of repetitive PMS intensity were assessed using conditions applied in random order across three sessions on separate days. A crossover design was employed, and repetitive PMS was delivered at three stimulus intensities: 0.9×MT (below the MT), 1.2×MT [(same as Nito et al. (2021) (13)], and a high intensity can induce maximal dorsiflexion movement of the wrist (high-intensity). The stimulus intensity required to induce maximal dorsiflexion ranged from 1.4 to 2.4×MT (mean ± SD: 1.8 ± 0.3×MT). The MT for the 0.9×MT, 1.2×MT, and high-intensity were 18.8 ± 2.9, 19.4 ± 2.1, and 20.1 ± 3.2, respectively (F_2,36_ = 2.513, p = 0.10, 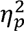= 0.123). A minimum one-day interval separated sessions to minimize carry-over effects. The intervention comprised five sessions, and a single session of the repetitive PMS was delivered for 2 s ON and 2 s OFF cycle for 3 min. MEPs were measured 3 min before intervention (baseline), just before repetitive PMS (T0), and after one session (T3), two sessions (T6), three sessions (T9), four sessions (T12), and five sessions of repetitive PMS (T15). The averaged MEP amplitude at each time point was normalized to the baseline and expressed as a percentage of the baseline value. M-max was also measured at T0 and T15.

#### Experiment 2: Persistence of facilitation after short, high-intensity repetitive PMS

To investigate the lasting effect of repetitive PMS, the stimulation was administered in three consecutive sessions, totaling 9 min, using the same stimulation cycle and frequency as in Experiment 1. The stimulus intensity was set to induce maximal dorsiflexion of the wrist (high-intensity). MEPs were measured 5 min before repetitive PMS (baseline), just before the repetitive PMS (Pre), and every 10 min for 50 min after the repetitive PMS (P0, P10, P20, P30, P40, and P50). The averaged MEP amplitude at each time point was normalized to the baseline and expressed as a percentage of the baseline value.

#### Experiment 3: Ischemia to probe the contribution of large-diameter afferents

To investigate the contribution of Ia afferent input to MEP enhancement, we applied repetitive PMS during ischemia and examined its effects on MEP amplitude. A blood pressure cuff placed around the participant’s upper arm was inflated to 220 mmHg pressure (32). After the inflation onset, the H-reflexes were elicited every 4 s by electrical stimulation to the radial nerve trunk, and the stimulus intensity was set to elicit the maximal peak-to-peak amplitude of the H-reflex. The H-reflex, which was absent at rest, was elicited during weak isometric wrist dorsiflexion. After the peak-to-peak amplitude of the H-reflex decreased to less than 10% of its preischemic size (6.4% ± 2.6% M-max; 12–20 min from the onset of ischemia), repetitive PMS was performed during ischemia in three consecutive sessions, totaling 9 min. The stimulus intensity was set to induce maximal dorsiflexion of the wrist (high-intensity). MEPs were measured 5 min before repetitive PMS (baseline), just before the repetitive PMS (Pre), and every 10 min for 50 min after the repetitive PMS (P0, P10, P20, P30, P40, and P50). The MEP measurement at P0 was performed after confirming that M-max remained unchanged compared to its size before ischemia (before ischemia, 5.9 ± 2.6 mV; after ischemia, 6.2 ± 2.0 mV; p = 0.79, d = 0.126).

#### Experiment 4: Estimation of afferent-driven motoneuron facilitation using MU recordings

To compare sensory input magnitude across PMS intensities, conditioning stimulation effects on a single MU were investigated (33). This approach is based on the premise that group Ia afferents have a monosynaptic connection to almost all homonymous motoneurons (34), enabling us to estimate the level of Ia afferent input by analyzing MU firings. Poststimulus time histograms (PSTHs) (bin width 0.2 ms or 0.5 ms) of the discharge of a voluntarily activated MU (approximately 100-ms firing interval) were constructed for the period ranging from 15 to 50□ms after conditioning stimulation. ECR MU discharges were recorded during isometric wrist extension at <5% maximal voluntary contraction. MU discharges were detected by the computer at intervals of approximately every 0.7 s, triggering the stimulator. The conditioning stimulation was triggered with a delay of about 70□ms after voluntary MU activation. The delay was set at a time that easily affected the next MU firing. In other words, the delay was set so that the afferent volleys by the conditioning stimulation would arrive at the motoneuron around the latest period of hyperpolarization due to the previous MU firing and just before voluntarily driven discharge of the motoneuron. Additionally, a firing-probability histogram was constructed under a no-stimulation control condition. The control and stimulated situations were alternated randomly (the same number of triggers) within a sequence. Each sequence generally comprised 200–800 stimulated and control situations. Conditioning stimulation effects were obtained by subtracting control-condition trigger counts from poststimulation values in each bin. Histogram subtraction was used to generate a cumulative sum curve confirming facilitation (35).

Conditioning PMS was applied to the radial nerve innervating the ECR. PSTHs were constructed using stimulation intensities of 0.9, 1.2, 1.5, and 1.8× MT to investigate PMS intensity effects on the firing probability of ECR MUs. For each single MU, PSTHs were constructed for each stimulation intensity, with the order of intensities randomized across measurements.

### Statistical analysis

The Shapiro–Wilk test was used to confirm that the normalized MEPs and M-max followed a normal distribution. In Experiment 1, a two-way repeated-measures mixed model analysis of variance (ANOVA) was used with INTENSITY (0.9×MT, 1.2×MT, and high-intensity) and TIME (T0, T3, T6, T9, T12, T15) as factors to compare normalized MEPs. When a significant interaction or main effects were determined, a two-way repeated-measures mixed model ANOVA was similarly used with INTENSITY (0.9×MT, 1.2×MT, and high-intensity) and TIME (T0 and T15) as factors to compare M-max.

Experiment 2 assessed lasting repetitive PMS effects on MEPs using a repeated-measures mixed model ANOVA with TIME as the main factor (Pre, P0, P10, P20, P30, P40, and P50).

Owing to the small sample size (n = 5), Experiment 3 reports individual data without statistical analysis.

In Experiment 4, effect latency and duration were defined as consecutive bins showing increased firing probability and were determined from fluctuations in cumulative sum curves by visual inspection from at least two investigators. Differences between poststimulation and control firing probabilities across consecutive-bin time windows were assessed using χ^2^-test. A repeated-measures mixed model ANOVA with main factor INTENSITY (0.9×MT, 1.2×MT, 1.5×MT, and high-intensity) was used to compare peak amplitudes derived from cumulative difference histograms. An unpaired t-test was performed to compare the peak duration induced by PMS and electrical stimulation.

Statistical significance was set at p < 0.05 for all comparisons. When significant main effects or interactions were found, post-hoc comparisons were performed using a paired t-test with Bonferroni adjustments. Effect sizes were reported as partial η² (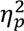) for ANOVA and Cohen’s d for the t-test. All statistical analyses, except the χ^2^-test, were performed using SPSS 30 (IBM, Armonk, NY, USA). The χ^2^-test was conducted using a PSTH analysis program (MTS0014, Gigatex, Japan). Group data are presented as mean ± SD in the text.

## RESULTS

### Experiment 1: Intensity-dependent effects of repetitive PMS on MEPs, M-waves

Figure 1 shows the time courses of the normalized MEPs. Notably, ECR MEP amplitudes were differentially modulated by stimulation intensity (Figure 1A). Two-way repeated-measures mixed model ANOVA revealed a significant main effect of INTENSITY (F_2,36_ = 11.524, p < 0.001, 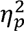 = 0.192) and TIME (F_5,90_ = 13.312, p < 0.001, 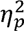= 0.470), and their interaction (F_10,180_ = 2.688, p = 0.004, 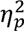 = 0.167) on the MEP amplitudes of the ECR. Time-factor comparisons revealed no changes following repetitive PMS at 0.9×MT, even after a total 15-min intervention (F_5,90_ = 0.885, p = 0.494, 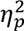 = 0.047). In contrast, repetitive PMS at 1.2×MT increased MEPs at T15 compared to T0 (p < 0.001, d = 1.047), whereas repetitive PMS at high-intensity induced an earlier facilitation, with MEP amplitudes increasing at T9 (p = 0.002, d = 0.997), T12 (p < 0.001, d = 1.585), and T15 compared to T0 (p < 0.001, d = 1.240). Comparison of the MEPs of ECR among different conditions showed a significant difference between 0.9×MT and high intensity repetitive PMS at T12 (p = 0.013, d = 0.859) and T15 (p = 0.002, d = 0.800), but no significant difference was observed in other combinations (p > 0.05).

**Figure 1.**
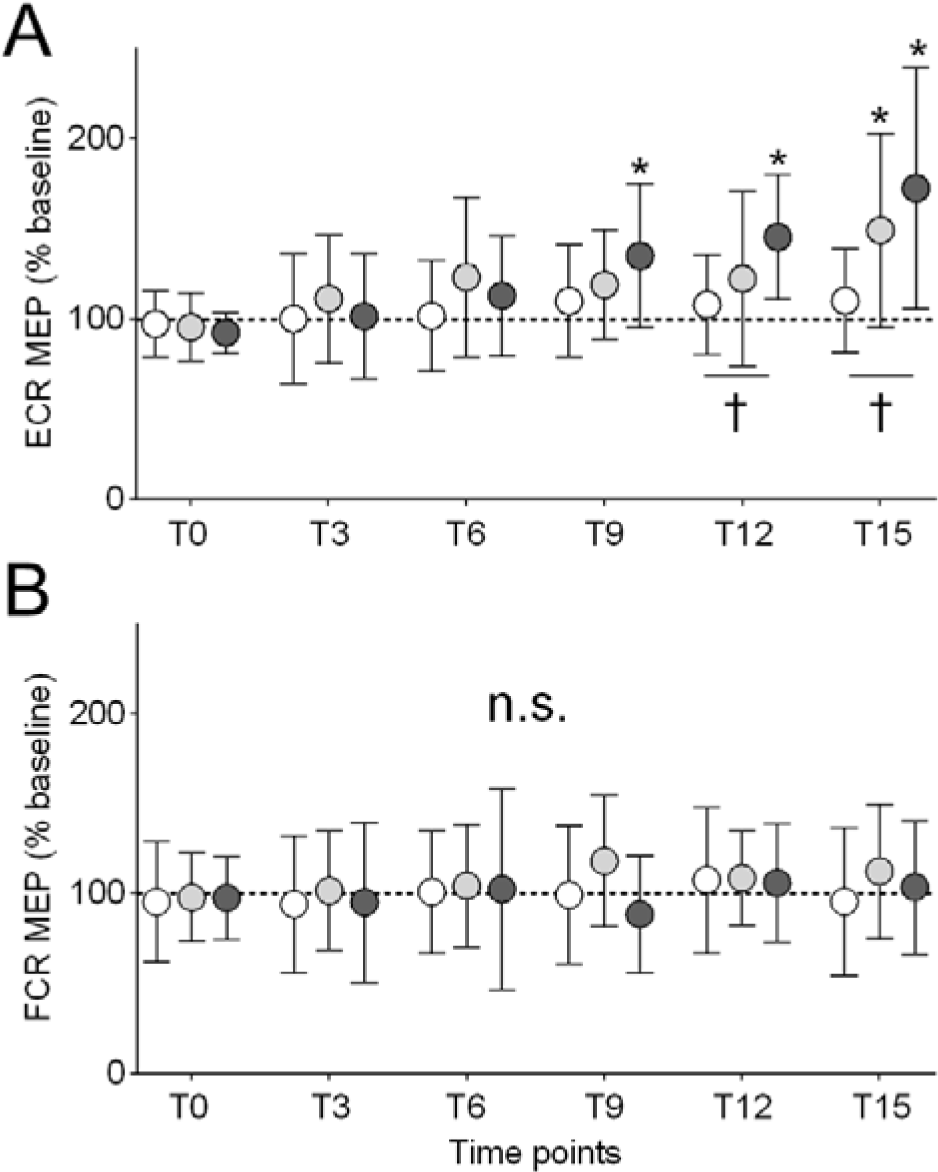
Effects of different intensities of repetitive peripheral magnetic stimulation (PMS) on motor-evoked potentials (MEPs). Changes in MEP amplitudes of the extensor carpi radialis (ECR; A) and flexor carpi radialis (FCR; B) muscles were recorded from 19 participants. The repetitive PMS was delivered at three intensities: below motor threshold (0.9×MT; white circle), above motor threshold (1.2×MT; light gray circle), and the intensity inducing maximum wrist dorsiflexion (high-intensity, dark gray circle). The ordinate shows MEP amplitude normalized to the baseline value for each participant, and the abscissa shows the time point of measurements: before repetitive PMS (T0), after one session (T3), two sessions (T6), three sessions (T9), four sessions (T12), and five sessions of repetitive PMS (T15). Each circle and error bar represent the mean value and SD, respectively. Asterisks indicate significant differences compared with “T0,” and the daggers indicate significant differences compared with the other intensities (p < 0.05).

FCR MEP amplitudes remained unaltered under any of these conditions (Figure 1B). A two-way repeated-measures mixed model ANOVA revealed no significant main effects of INTENSITY (F_2,36_ = 0.594, p = 0.104, 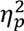 = 0.033) or TIME (F_5,90_ = 0.820, p = 0.536, 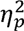 = 0.062), and their interaction (F_10,180_ = 0.968, p = 0.473, 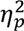 = 0.051) on the FCR MEP amplitudes.

The M-max amplitudes remained unchanged under any of these conditions. The mean values of M-max at T0 and T15 were 7.9 ± 3.2 mV and 8.2 ± 3.2 mV for 0.9×MT repetitive PMS, 7.5 ± 2.6 mV and 7.5 ± 2.8 mV for 1.2×MT repetitive PMS, 7.3 ± 2.4 mV and 7.7 ± 2.7 mV for high-intensity repetitive PMS, respectively. A two-way repeated-measures mixed model ANOVA revealed no significant main effects of INTENSITY (F_2,36_ = 1.242, p = 0.301, 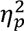 = 0.065) or TIME (F_1,18_ = 2.065, p = 0.168, 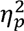 = 0.103), and their interaction (F_2,36_ = 0.432, p = 0.652, 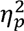 = 0.023).

### Experiment 2: Persistence of facilitation after short, high-intensity repetitive PMS

Figure 2 shows the changes in MEPs of ECR before and after 9 min of repetitive PMS. Repetitive PMS markedly increased MEPs in stimulated ECR, whereas no significant changes were observed in FCR. A repeated-measures mixed model ANOVA for ECR MEPs revealed a significant main effect of Time (F_6,96_ = 5.153, p < 0.001, 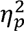 = 0.244). Post-hoc tests revealed that ECR MEPs were significantly increased at P0 (p < 0.001, d = 1.174), P10 (p < 0.001, d = 1.484), P20 (p = 0.024, d = 0.968), and P30 (p = 0.037, d = 0.725) compared to Pre values, whereas no significant changes were observed at other time points (p > 0.05). In contrast, a repeated-measures mixed model ANOVA for FCR MEPs revealed no significant main effect of Time (F_6,96_ = 0.985, p = 0.440, 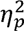 = 0.059).

**Figure 2.**
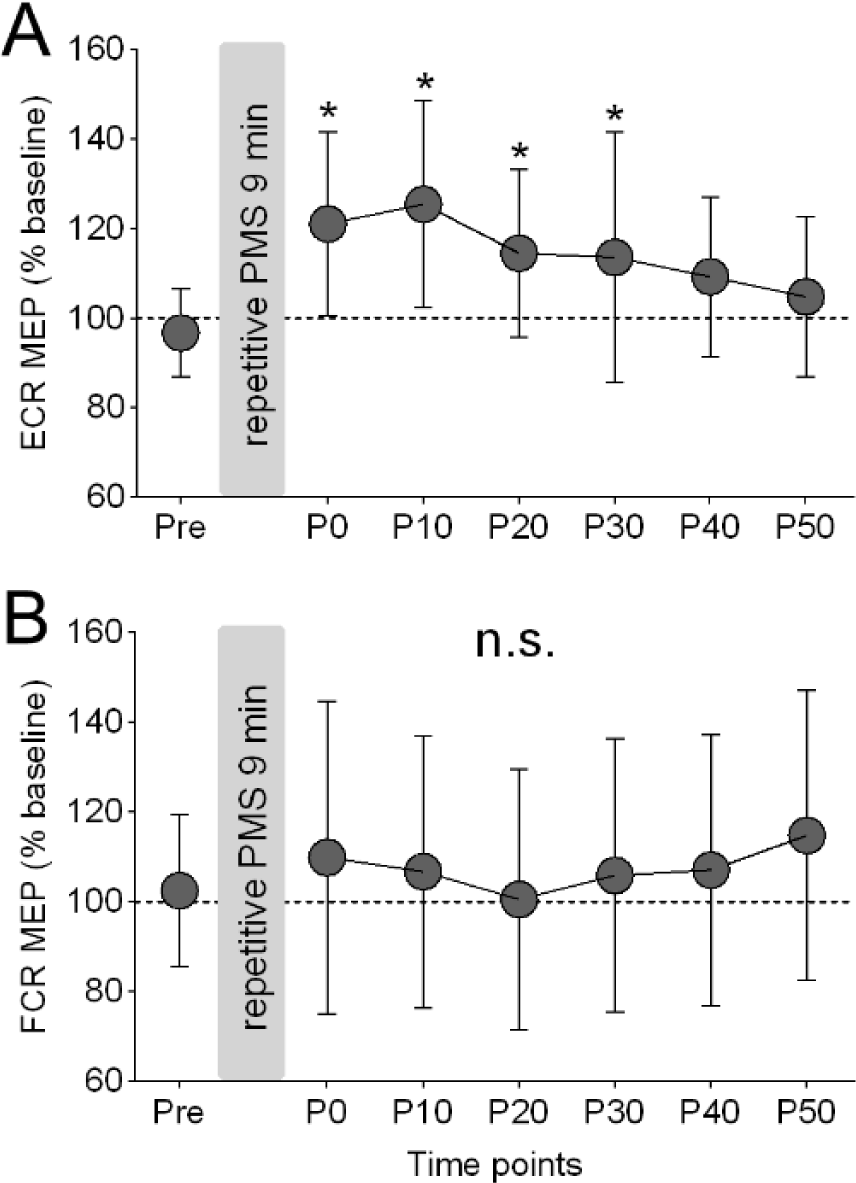
Lasting effects of repetitive PMS on MEPs. Time courses of MEP changes in the ECR (A) and FCR (B) muscles were recorded from 17 participants. The ordinates show MEP amplitude normalized to baseline amplitude for each participant, and the abscissae show the time point of measurements: before (Pre) and after the repetitive PMS (P0–P50). Each circle and error bar represent the mean value and SD, respectively. Asterisks indicate significant differences compared with “Pre.”

### Experiment 3: Ischemia to probe the contribution of large-diameter afferents

Figure 3 illustrates the individual changes in ECR MEPs for five participants who participated in Experiments 2 and 3. As shown in Figure 3A, which indicates the individual data underlying Figure 2, a robust increase in MEPs was confirmed. Conversely, repetitive PMS applied after ischemic blockade of Ia afferent input produced no observable MEP changes (Figure 3B), suggesting a contribution of Ia afferent input to repetitive PMS–induced MEP augmentation.

**Figure 3.**
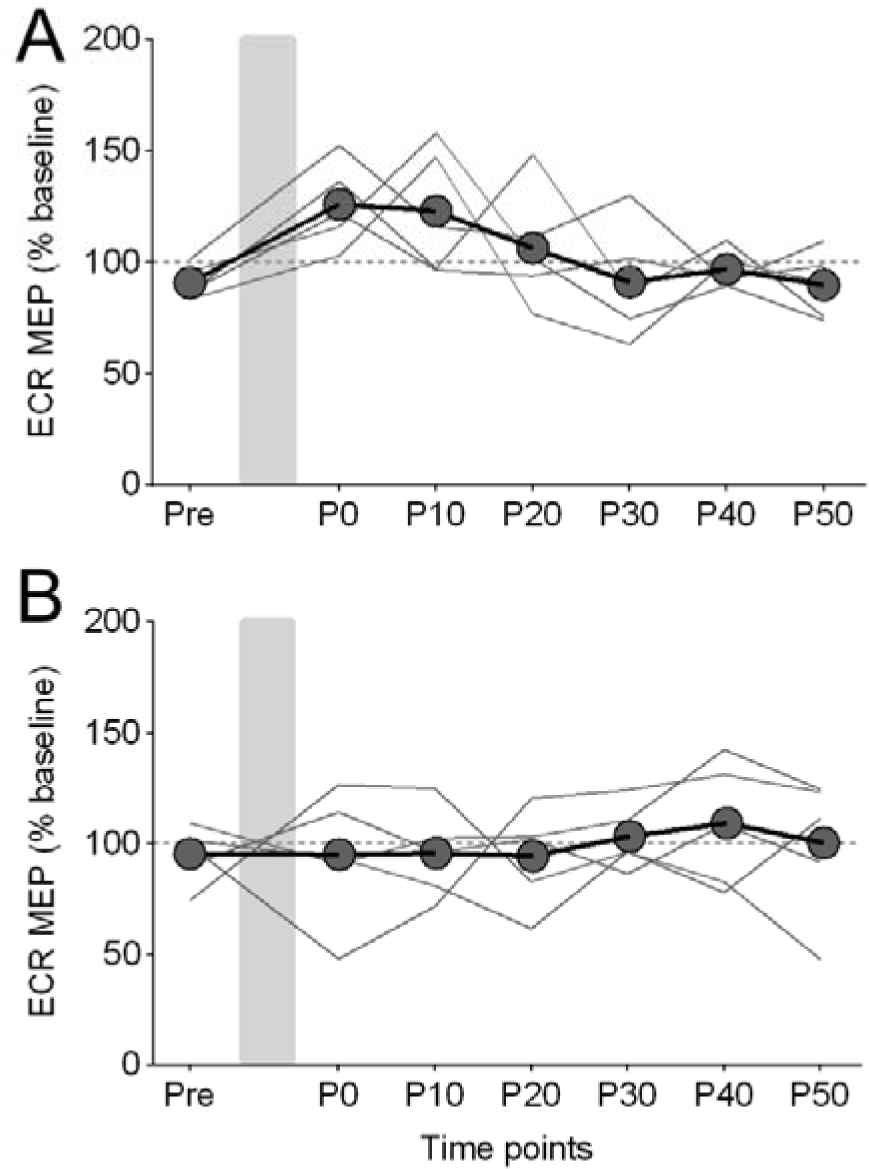
Effects of ischemia on MEP facilitation induced by repetitive PMS. The time courses of MEP changes in the ECR were recorded from five participants without (A) and with ischemia (B). Individual MEP changes and their mean values are represented as gray lines and circles, respectively. The ordinate and abscissa are the same as those in Figure 2.

### Experiment 4: Estimation of afferent-driven motoneuron facilitation using MU recordings

Figure 4 shows a representative facilitation induced by PMS with various intensities. A significant peak (representing an increase in the firing probability) of PSTH was observed 22.5–24.0 ms after PMS using 1.8×MT (p□<□0.001). In this MU, a peak was observed at the same latency across all stimulus intensities from 1.2×MT to 1.8×MT, whereas the peak size, as indicated by the cumulative sum curve, decreased upon PMS intensity reduction. The PMS at 0.9×MT did not affect the MU firing probability (p > 0.05). A total of 27 MUs from eight participants were studied. Because stable MU firing could not be maintained over prolonged periods, the number of analyzed MUs varied slightly across stimulation intensities. A significant peak was observed in all 25 MUs when PMS was applied at 1.8×MT (100%), in 17 of 24 MUs at 1.5×MT (71%), and in 14 of 25 MUs at 1.2×MT (56%). In contrast, no significant peak was detected in any of the 25 MUs using PMS at 0.9×MT (0%) (Figure 4B). Furthermore, the peak amplitudes obtained from the cumulative sums of the difference histograms increased progressively with stimulation intensity (Figure 4C). A repeated-measures mixed model ANOVA revealed a significant main effect of INTENSITY (F_3,66.065_=23.936, p<0.001, 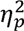=0.486). These findings suggest that higher PMS intensities are associated with greater afferent input, whereas PMS delivered below the MT does not elicit afferent input.

**Figure 4.**
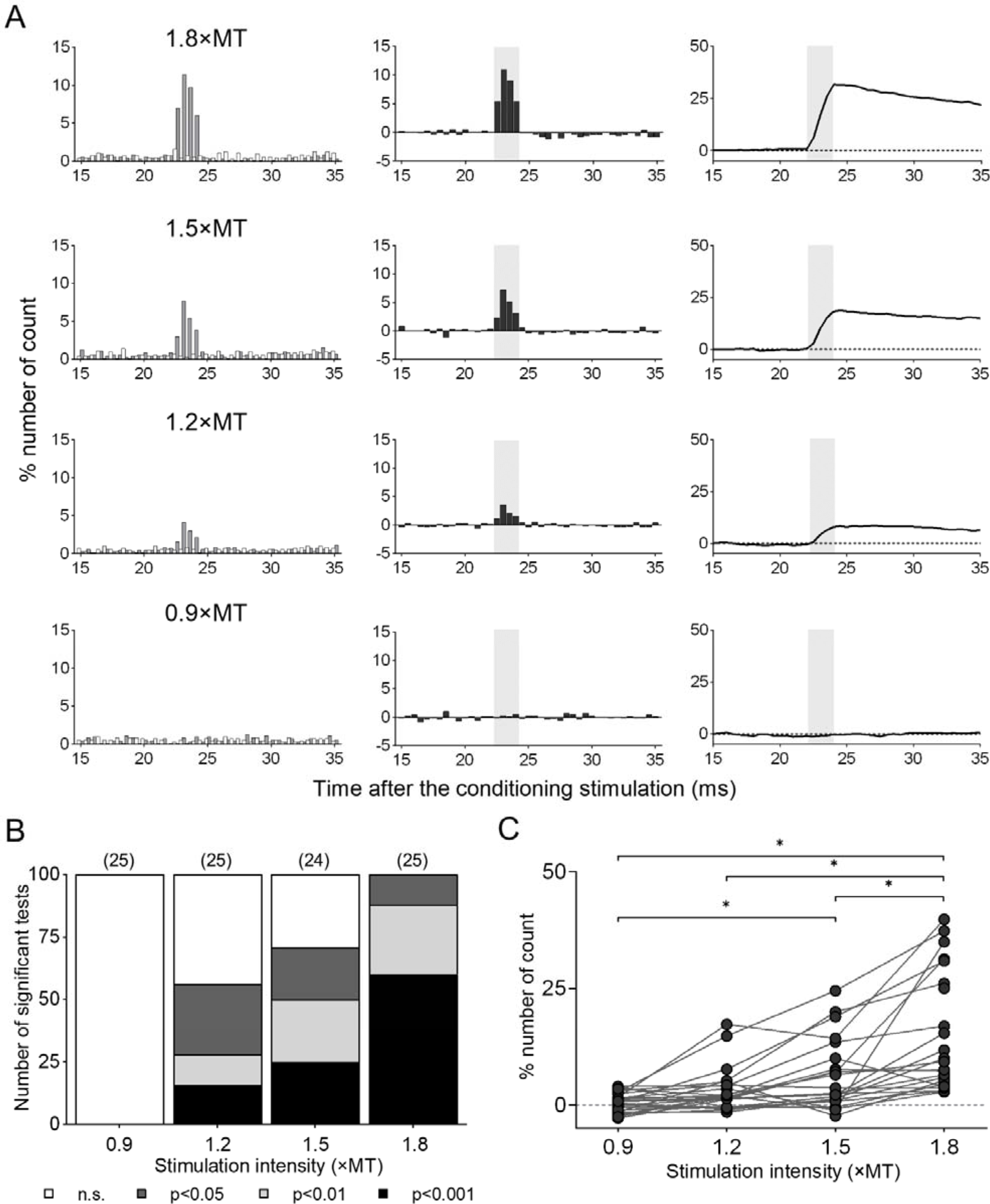
Changes in the firing probability of an ECR motor unit following different intensities of PMS. A, Left: Time histograms were obtained without PMS (white bar) or with PMS to the radial nerve innervating the ECR (black bar) within a sequence. Each histogram was constructed from 600 triggers. Middle: Each column (dark gray bar) represents the difference between the situations with and without PMS. The analysis time window (light gray area) was determined from several consecutive bins of the time histograms obtained during 1.8×MT PMS. Right: Cumulative sums are obtained from each subtracted histogram. The ordinates represent the number of counts as a percentage of the number of triggers, and the abscissae represent the latency after stimulation. Bin width: 0.5 ms. B, Stacked bar graph showing the proportion of motor units that exhibited significant peaks in the time histograms under each stimulation intensity. The numbers in parentheses indicate the number of motor units. C, Changes in the peak amplitude obtained by cumulative sums within the analysis time window of the difference histograms under different stimulation intensities. Individual changes and their mean values are represented as gray lines and circles, respectively. Asterisks indicate significant differences revealed by post-hoc tests (p < 0.05).

Figure 5 shows an example of the effects of PMS and electrical stimulation of the radial nerve trunk on single MU firings. The latency of the peak was 22.0 ms for PMS and 20.4 ms for electrical stimulation, and the duration was 2.4 ms and 2.4 ms, respectively. A total of 15 MUs from 8 participants were studied. The difference latency of the peaks was 21.4 ± 2.0 ms for PMS and 20.4 ± 2.0 ms for electrical stimulation, respectively, indicating that the electrical stimulation–induced peak latency was 1.0 ± 0.7 ms shorter than that of PMS. Considering the distance between the two stimulation sites (8.2 ± 3.4 cm), the central latency of the PMS-induced peak can be considered equivalent to that of monosynaptic Ia facilitation induced by electrical stimulation (36). The peak duration was 2.4 ± 1.0 ms for PMS and 2.2 ± 0.8 ms for electrical stimulation, respectively, indicating no significant difference between the two conditions (p = 0.402 using unpaired t-test, d = 0.157). These findings suggest that PMS activates Ia afferents and generates excitatory postsynaptic potentials (EPSPs), with the initial component mediated by a monosynaptic pathway.

**Figure 5.**
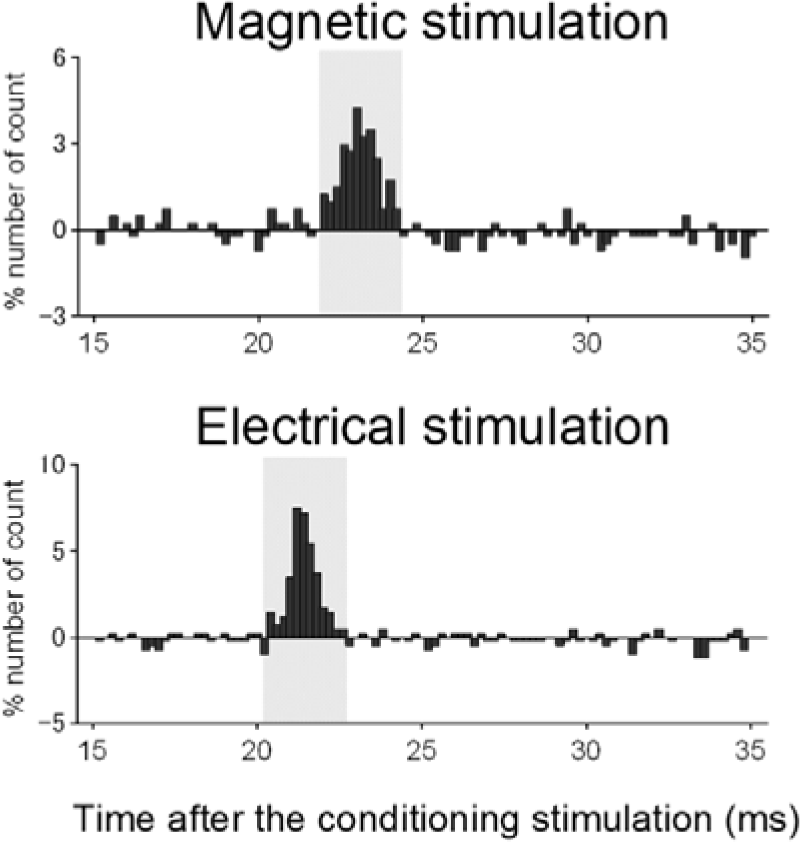
Changes in the firing probability of an ECR motor unit following electrical and magnetic stimulation. Time histograms were obtained by subtracting the situation without stimulation from that with stimulation. Each histogram was constructed from 400 triggers. The ordinate and abscissa are the same as in Figure 4A. Bin width: 0.2 ms. The difference between the latencies of peaks induced by electrical stimulation (20.4 ms) and magnetic stimulation (22.0 ms) was 1.6 ms. The distance between the two stimulation sites was 10 cm.

## DISCUSSION

The present study aimed to clarify how varying intensities of repetitive PMS applied to wrist extensor muscles influence CSE. Our results yielded three main findings. First, repetitive PMS facilitated CSE in an intensity-dependent manner, with high-intensity stimulation inducing earlier facilitation than 1.2×MT, while 0.9×MT produced no effect. Second, the facilitation was muscle-specific and not attributable to peripheral changes, as FCR MEPs and ECR M-max remained unchanged. Third, the abolition of facilitation during ischemia together with a short-latency peak in MU firing via a monosynaptic path supports a major contribution of large-diameter muscle afferents, consistent with a substantial Ia component, to PMS-induced corticospinal facilitation.

### Group I Afferents Responsible for CSE Enhancement

Group I afferents from wrist extensor muscles are most likely the afferent fibers responsible for CSE enhancement. In the present study, we demonstrated that a suprathreshold PMS increased MEPs. Specifically, high-intensity PMS-induced increased MEPs, achieving this effect only after a short duration of intervention. Furthermore, MU firing analysis showed similar PSTH peak latencies across suprathreshold PMS intensities, whereas peak amplitudes increased with stimulation intensity. Additionally, the PSTH duration was comparable to that induced by electrical stimulation to the radial nerve trunk, which is known to elicit a monosynaptic component mediated by Ia afferents (37, 38), regardless of the stimulation intensity of PMS. The peak amplitude and duration of the PSTH are considered to reflect the size and rise time of the composite EPSPs, respectively (39, 40). Higher PMS intensities may have recruited a greater number of Ia afferents, thereby eliciting larger EPSPs. Consequently, stronger intensities of PMS may have induced increased CSE within a shorter intervention period compared to weaker intensities. Ischemic nerve block is known to preferentially attenuate large-diameter afferents (32). Accordingly, the absence of MEP facilitation during ischemia supports a major contribution of large-diameter muscle afferents (i.e., group I pathways) to PMS-induced corticospinal facilitation. Although ischemia does not isolate Ia afferents exclusively, the monosynaptic timing and narrow duration of MU facilitation are compatible with a substantial Ia component. Because repetitive PMS was delivered after the H-reflex was reduced to <10% of baseline, these findings collectively suggest that attenuation of large-diameter afferent input abolished the facilitatory effect on CSE.

PMS at subthreshold intensities did not alter CSE or MU firings in the present study. This interpretation is supported by a previous report by Panizza et al. (1992) (29), which demonstrated two key findings regarding PMS: First, sensory fibers exhibit a higher threshold than motor fibers. Second, the H-reflex—elicited by monosynaptic excitation from the homonymous group Ia afferents—is not induced at intensities lower than the MT. Additionally, previous studies on electrical stimulation have shown contrasting effects: suprathreshold electrical stimulation of the median nerve trunk at the wrist increases MEPs, whereas subthreshold stimulation decreases MEPs (26–28). Furthermore, cutaneous nerve stimulation of the index finger originating from the median nerve also decreases MEPs (26). Comparable conduction velocities between the fastest cutaneous and group Ia afferents (41) suggest similar recruitment thresholds. Therefore, the inhibitory effect of subthreshold stimulation on CSE may result from the preferential activation of cutaneous afferents rather than of group Ia afferents. In contrast, PMS may be more effective in increasing CSE by preferentially activating Ia afferents and providing stronger ascending proprioceptive input. Although PMS was applied transcutaneously and may activate cutaneous afferents beneath the stimulation site, magnetically induced eddy currents directly stimulate deeper tissues and recruit superficial cutaneous afferents less effectively than electrical stimulation (8). Thus, cutaneous afferent contributions at the stimulation site are likely smaller than those typically recruited by peripheral electrical stimulation, although this study did not directly quantify such observation. Therefore, these findings suggest that subthreshold PMS is insufficient to excite the sensory afferents necessary to modulate CSE, thereby failing to induce either enhancement or suppression of MEPs.

### Changes in CSE Induced by Repetitive PMS

Previous studies (7, 12, 13) have revealed that repetitive PMS at suprathreshold intensity increases MEPs. Moreover, reductions in short-latency intracortical inhibition and increases in intracortical facilitation have been reported following repetitive PMS (12, 13). In contrast, MEPs evoked by transmastoid electrical stimulation were not altered by repetitive PMS (Omiya et al., *under review*), which is thought to reflect changes in the efficiency of synaptic transmission between the cortex and spinal motor neurons or the excitability of spinal motor neurons themselves (42, 43). Furthermore, the excitability of the spinal reflex loop in the muscle receiving PMS appears to remain unchanged, as demonstrated using the H-reflex (13). In addition, M-max remained unchanged following repetitive PMS, indicating preserved neuromuscular transmission efficacy. Overall, these findings suggest that augmented MEPs are predominantly attributable to changes at supraspinal levels (e.g., cortical and/or corticospinal synaptic efficacy), although the present design does not fully exclude contributions from spinal mechanisms.

Prior work demonstrated sustained MEP enhancement for up to 60 min after 15 min of repetitive PMS at 1.2×MT (13). In contrast, the present study demonstrated that higher intensities of PMS-induced greater increases in MEPs following only 9 min of the intervention, but the effect persisted for only 30 min. These findings suggest a trade-off between the intensity and duration of PMS-induced effects: higher intensities can induce greater increases in CSE within a shorter intervention time, but the effects are less sustained compared with those induced by lower intensities. Therefore, high-intensity PMS, capable of inducing rapid increases in CSE, may provide practical advantages in clinical settings with time constraints. A limitation of the present study is that the duration of CSE changes following high-intensity PMS applied over longer intervention periods (e.g., 15 min) was not investigated. It is conceivable that extending the duration of high-intensity stimulation may prolong CSE enhancement. Future studies are warranted to determine whether longer high-intensity PMS protocols can combine the advantages of rapid induction and sustained effects.

### Methodological Considerations

Despite these strengths, several methodological considerations warrant acknowledgment. First, the ischemic nerve block procedure in Experiment 3 was performed in a small sample (n = 5). This small sample size limits the generalizability of our findings that large-diameter afferent input—with a substantial Ia component—mainly contributes to PMS-induced facilitation. Second, despite marked ischemia-induced H-reflex suppression indicating reduced group Ia afferent input, potential contributions from other large-diameter afferents, including group Ib afferents, remain possible. These afferents may also have been partially affected, potentially contributing to the observed increase in MEPs (23–25). Third, repetitive PMS–induced coil heating constitutes a methodological limitation. At frequencies higher than 25 Hz or at higher stimulation intensities, continuous intervention may become challenging due to the rapid rise in coil temperature. Therefore, the parameters used in this study were selected to balance the induction of corticospinal facilitation with the physical capacities of the stimulator.

### Clinical Implications

Numerous studies have reported that repetitive PMS contributes to improved motor dysfunction following CNS lesions (7, 9–11, 16, 18–20, 22), suggesting promising therapeutic potential. Repetitive PMS–induced increases in CSE likely contribute, at least in part, to these functional improvements (7).

Beyond these potential benefits for functional recovery, PMS offers practical advantages for clinical application: PMS can directly stimulate deep tissues without penetrating the skin to induce eddy currents (8), and is therefore considered less likely to cause discomfort during stimulation. Indeed, none of the participants reported any discomfort during PMS, even at an intensity sufficient to elicit maximal wrist dorsiflexion, and no participant found it difficult to continue the experiment.

While the present findings provide evidence for the underlying mechanism of PMS, its clinical applicability requires further investigation. The present study examined only the immediate effects of repetitive PMS on wrist extensors in healthy individuals; whether similar facilitatory effects occur in individuals with CNS lesion–related motor dysfunction or are sustained across repeated, longer-term interventions remains to be determined. Furthermore, although we demonstrated that repetitive PMS increases CSE, the functional implications of this enhancement remain to be fully elucidated. Future studies incorporating behavioral or functional outcome measures should clarify whether PMS-induced increases in excitability translate into meaningful improvements in motor performance or clinical function.

## CONCLUSIONS

Repetitive PMS at intensities above the MT enhances CSE, likely mediated predominantly by large-diameter muscle afferents (group I afferents), consistent with a substantial Ia component. Short-duration interventions at higher stimulation intensities produce greater CSE enhancement than lower-intensity stimulation. In addition, stronger stimulation intensities are associated with greater afferent input. These findings suggest that group Ia afferents from muscle spindles markedly contribute to CSE modulation. These results provide insight into the mechanisms underlying PMS-induced neuroplasticity and may help optimize stimulation parameters for clinical applications in neurorehabilitation.

## LIST OF ABBREVIATIONS

CSE: corticospinal excitability
CNS: central nervous system
ECR: extensor carpi radialis
EMG: electromyography
EPSP: excitatory postsynaptic potential
FCR: flexor carpi radialis
MEP: motor-evoked potential
M-max: maximum direct motor response
MT: motor threshold
MU: motor unit
PMS: peripheral magnetic stimulation
PSTH: poststimulus time histogram
TMS: transcranial magnetic stimulation

## Data availability

The datasets supporting the conclusions of this article are available from the corresponding author on reasonable request.

## Acknowledgments

The authors thank all the participants who engaged in this study.

## Author contributions

M.N., N.M. and T.Y. conceived and designed research, K.Y., M.N., D.M., A.O., and K.S. performed experiments and analyzed data, K.Y., M.N., H.F., and T.Y. interpreted results of experiments, M.N. prepared figures, K.Y., M.N. and T.Y. drafted manuscript, K.Y., M.N., D.M., A.O., K.S., T.K., D.K., N.M., H.F., and T.Y. approved the final version of manuscript.

